# Microbial network rewiring mediates the function-biosafety trade-off in transboundary Pamir lakes

**DOI:** 10.64898/2026.05.19.726157

**Authors:** Yuzheng Gu, Zhaojing Liu, Congrui Liu, Xiaoyun gou, Yanhan Ji, Baozhan Wang, Xu Liu, Jiandong Jiang

## Abstract

The Pamir Plateau is a transboundary water tower whose source lakes serve as critical biogeochemical hubs with implications for downstream freshwater security. However, it remains unclear how environmental shifts in these high-altitude lakes reshape the microbial communities that drive ecosystem functioning and water safety. Here, we conducted a multi-omics survey across 20 lakes spanning Chinese and Tajikistani Pamir. Our results revealed that prokaryotes exhibited lower diversity but higher among-lake connectivity in China, while eukaryotes showed higher diversity but stronger dispersal limitation. These contrasting biogeographic responses triggered profound rewiring of microbial associations. Under intensified anthropogenic pressures, Chinese cross-kingdom networks decoupled from environmental constraints and became more centralized and complex. Conversely, Tajikistani lakes maintained more modular networks governed by hydrochemical filtering. Critically, this rewiring mediated a trade-off between multifunctionality and potential biosafety risk, with higher element cycling abundances in Chinese lakes, whereas Tajikistani lakes harbored larger biosafety burden dominated by virulence, pathogen, and toxic-algae potential. Incorporating network topology also substantially improved the prediction of these ecological consequences. These findings highlight the importance of network-informed monitoring and management strategies to safeguard ecosystem sustainability in transboundary Pamir lakes under global change.

## Introduction

The Pamir Plateau is a transboundary high-mountain water tower in arid Central Asia, spanning Tajikistan, Kyrgyzstan, Afghanistan, and China^1^. Snow- and glacier-fed runoff from this region contributes to major downstream dryland basins, including the Amu Darya and Tarim basins, where water availability is closely linked to irrigation, ecosystem stability and human livelihoods^2,3^. This water-tower system is increasingly affected by global changes and local human activities, including glacier retreat, altered runoff regimes, shifting precipitation patterns, atmospheric deposition, grazing, tourism, and contaminant inputs^1,2^. Within this system, high-altitude lakes occupy a pivotal position. They not only serve as natural reservoirs of headwater runoff but also as sensitive sentinels of environmental changes because they can integrate hydrological, chemical, and biological inputs from the surrounding landscape and atmosphere ^4–7^. These lakes are also crucial biogeochemical hubs, where nutrients and organic matter are retained, transformed, buried, or exported, thereby influencing primary productivity and nutrient cycling of lake ecosystems^8,9^. Meanwhile, contaminants and potential biological hazards accumulated or generated in these lakes can be transported downstream through lake outflows and connected river networks, with great implications for water security^8,10^. Therefore, the ecological conditions of Pamir lakes are directly relevant to ecosystem functioning and downstream water quality. However, the microbial communities that regulate these lake ecosystem properties remain poorly characterized on the Pamir Plateau.

Although low temperature, intense ultraviolet radiation, and generally oligotrophic conditions impose severe constraints^12^, these lakes can harbor diverse and active prokaryotic and eukaryotic microorganisms, which form a substantial component of local biodiversity^4,6,13^. These microorganisms play essential roles in driving element cycling (e.g., carbon fixation, nitrogen transformation, and sulfur mineralization) and regulating food-web dynamics^14–17^. Moreover, lake microbiomes may contain pathogens or harmful algal taxa, or carry antibiotic resistance genes (ARGs) and virulence factors (VFs), making them relevant to the assessment of biological safety in source waters^18,19^. Previous studies have largely examined high-altitude lake microbiomes through diversity patterns, taxonomic composition, and community assembly, revealing predictable shifts along environmental gradients such as salinity, pH, and temperature, and indicating that their assembly is strongly governed by deterministic processes^20–22^. However, this composition-centered perspective overlooks the connected organization of microbial communities. Recent ecological theory emphasizes that, in a rapidly changing world, what determines ecosystem function and stability is not only which species are present, but how their interactions are rewired in response to environmental change^23,24^. This network rewiring can occur even when species turnover is modest, and may provide a mechanism by which environmental perturbations are translated into ecosystem-level consequences^23,24^. Yet in high-altitude lake microbiomes, the topological dynamics of cross-kingdom co-occurrence networks, and critically how they translate biogeographic divergence into ecological consequences, are largely unknown.

Therefore, we conducted a large-scale field survey of 20 high-altitude lakes across the Pamir Plateau, spanning both Chinese and Tajikistani regions (Fig.1a). These two regions provide a natural contrast in environmental conditions across the transboundary Pamir Plateau. Tajikistani lakes in the western Pamir are more directly exposed to moisture-bearing westerlies, while Chinese lakes lie mainly on the leeward eastern side ^25,26^. This marked difference was also reflected in our sampled Chinese and Tajikistani lakes, in terms of climate regime, water physicochemical properties, and the intensity of human activities (Fig.1b). We therefore used the country-level grouping as an integrative proxy for regional environmental divergence, rather than as a political boundary per se. By jointly profiling prokaryotic and eukaryotic communities, cross-kingdom networks, and functional potentials between Chinese and Tajikistani lakes, we aimed to: (1) characterize how regional environmental divergence shapes the biogeography and assembly of prokaryotic and eukaryotic communities; (2) determine how cross-kingdom microbial networks rewire between regions and identify their drivers; and (3) evaluate whether and how shifts in networks affect microbial functions and biosafety risk. Filling these knowledge gaps is essential for establishing a mechanistic basis for the conservation, risk assessment and ecological restoration of high-altitude lake ecosystems under global change.

## Results and Discussion

### Lake microbial biogeography on the Pamir Plateau

Prokaryotic and eukaryotic communities exhibited divergent biogeographic responses across the Pamir Plateau. Prokaryotic alpha diversity, including richness, Shannon, and Simpson indices, was significantly higher in Tajikistani lakes. By contrast, eukaryotic richness did not differ between regions, but eukaryotic Shannon and Simpson indices were significantly higher in Chinese lakes (Fig. 1c and Supplementary Fig. 1), indicating a regional shift in evenness rather than richness alone. Principal coordinate analysis (PCoA) revealed significant geographic segregation for both prokaryotic and eukaryotic communities (Fig. 1d). At the phylum level, Proteobacteria and Actinobacteriota dominated prokaryotic communities in both regions, but Chinese lakes hosted a notable expansion of Deinococcota, whereas Bacteroidota was more prevalent in Tajikistan (Supplementary Fig. 2). Given the well-known resistance of Deinococcus-related lineages to radiation, desiccation, and oxidative stress, this enrichment provides a taxonomic signal with stronger selection for stress-tolerant prokaryotic taxa in Chinese region^27^. Among eukaryotes, Chlorophyta and Ochrophyta constituted a larger proportion in Tajikistan, while Basidiomycota and Ascomycota were more abundant in China (Supplementary Fig. 2). Importantly, this regional divergence extended deep into the phylogeny (Fig. 1e). For example, within the class Gammaproteobacteria, the order Burkholderiales was enriched in Tajikistani lakes, while the order Pseudomonadales exhibited distinct intra-clade segregation, with the genus Acinetobacter lineages proliferating in China and the genus Pseudomonas lineages being enriched in Tajikistan. Such phylogenetic sorting suggests that regional environmental divergence acted not only by replacing major taxonomic groups, but also by selecting among closely related lineages with different ecological strategies.

**Figure 1.**
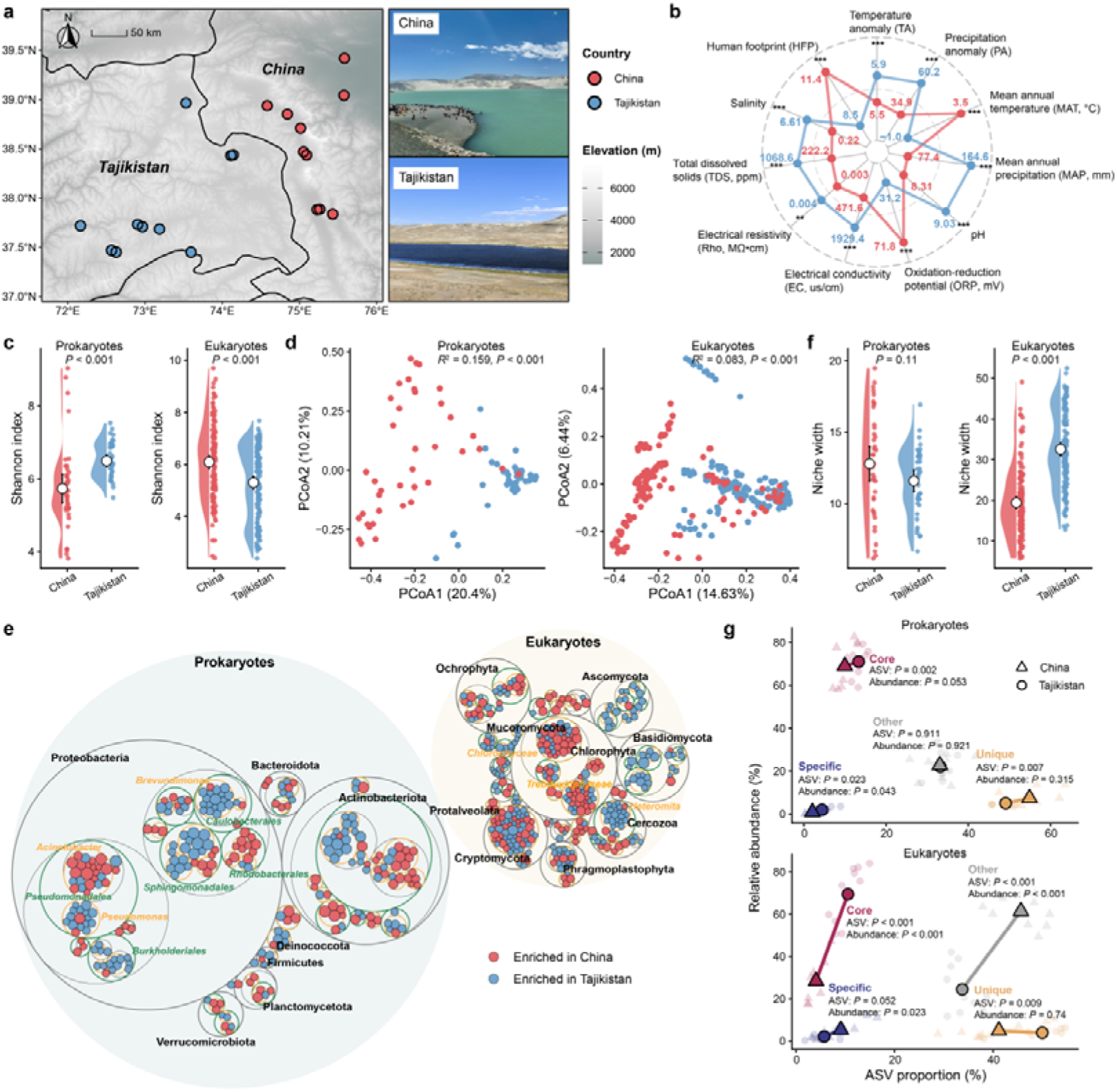
Geographic distribution and compositional divergence of lake microbiomes across the Pamir Plateau. **a**, Location of the sampling sites, which include ten lakes each in Chinese and Tajikistani regions. **b**, Differences of environmental variables between regions, including temperature anomaly, precipitation anomaly, mean annual temperature, mean annual precipitation, pH, oxidation-reduction potential, electrical conductivity, electrical resistivity, total dissolved solids, salinity, and human footprint. Asterisks indicate statistical significance (\*\*\**P* < 0.001, \*\**P* < 0.01, and \**P* < 0.05). **c**, Shannon index of prokaryotic and eukaryotic communities. White circles and error bars indicate the mean ± se. **d**, PCoA for prokaryotic and eukaryotic communities with PERMANOVA. **e**, Differences at the ASV level. Red and blue bubbles represent ASVs with significant enrichment (FDR < 0.05) in Chinese and Tajikistani regions, respectively, based on statistical analysis with ALDEx2. Bubble sizes represent the ALDEx2 effect size between the relative abundance of ASVs in China and Tajikistan. The largest circles in black represent the phylum level, the circles in green represent the order level, and the circles in orange represent the genus level. **f**, Niche width of prokaryotic and eukaryotic communities. White circles and error bars indicate the mean ± SE. **g**, Shifts in ASV proportion and relative abundance of core, specific, unique, and other sub-communities between Chinese and Tajikistani regions. *P* values were calculated using two-sided Wilcoxon rank-sum tests.

In addition, prokaryotes displayed comparable niche widths across both regions (Fig. 1f). Conversely, eukaryotes in Tajikistan showed significantly broader niches than in China. We further divided the communities into core and peripheral (specific, unique, and other) sub-communities to clarify these differences (Fig. 1g). Prokaryotes maintained a classic abundance-occupancy structure in both regions, where a small fraction of widespread core amplicon sequence variant (ASVs) accounted for most of the relative abundance. This pattern is consistent with aquatic microbial studies showing that high-occupancy and high-abundance taxa often represent a resident core shaped by local environmental sorting, whereas low-occupancy taxa are more likely to reflect intermittent recruitment or local restriction^28,29^. Therefore, although prokaryotic ASV proportions differed between regions, their stable abundance suggests that the prokaryotic communities retained a conserved structural backbone across the Pamir Plateau. Strikingly, the relative abundance of core eukaryotes in Chinese lakes sharply dropped by approximately 30% compared to their dominance in Tajikistan (Fig. 1g). Instead, other taxa expanded to fill this gap, highlighting that Chinese eukaryotes may have shifted to an opportunist-driven fragmented structure, which is consistent with their narrower niche widths.

We then used source tracking to evaluate community connectivity between lakes, with higher source contributions indicating stronger compositional exchange or similarity among lake communities (Supplementary Fig. 3a). Prokaryotic connectivity was significantly higher among Chinese lakes, whereas eukaryotic connectivity reached higher levels in Tajikistani lakes. This could be due to the facts that lower human disturbance and frequent waterbird activity in Tajikistan may facilitate the movement of eukaryotic propagules among lakes^30,31^, which aligned with the weaker eukaryotic distance-decay relationship (DDR) observed in this region (Supplementary Fig. 3b). In contrast, Chinese lakes showed a complete breakdown of prokaryotic DDR, which could be attributed to the stronger human footprint here, such as tourism, road access, and runoff inputs, further enhancing the spread of prokaryotes among lakes. This is because prokaryotic microorganisms are generally smaller and more easily passively transported than eukaryotes^32,33^. Null-model analyses of community assembly provided a mechanistic basis for these contrasts (Supplementary Fig. 3c). Specifically, prokaryotic assembly in China was largely driven by stronger deterministic processes alongside weaker dispersal limitation. By contrast, eukaryotes in China were severely constrained by dispersal limitation, whereas Tajikistani eukaryotes were structured predominantly by homogeneous selection. These results revealed that the geographic barriers impose distinct and even contrasting impacts on prokaryotic versus eukaryotic kingdoms. Procrustes analysis further showed significant cross-kingdom coupling in both regions, which was stronger in Tajikistan than in China (Supplementary Fig. 4). Together, these results imply that regional environmental divergence reshaped microbial community composition and ecological strategy through deterministic selection and dispersal, thereby disrupting the biogeographic coordination between prokaryotic and eukaryotic assemblages.

### Microbial co-occurrence networks

To further evaluate the alteration of possible ecological interactions among prokaryotic and eukaryotic members across regions, we constructed cross-kingdom co-occurrence networks (Fig. 2a). Our results showed that the overall network diverged between Chinese and Tajikistani lakes (dissimilarity = 0.916), which was driven by rewiring (62.8%) rather than node turnover (37.2%) (Fig. 2b), suggesting that regional divergence reorganized microbial association patterns even when many taxa were retained across regions^23^. While prokaryotes and eukaryotes shared comparable topological importance in Tajikistan, eukaryotic nodes became topologically dominant in China, exhibiting a nearly two-fold increase in their average degree relative to prokaryotes (Fig. 2c). This resulted in a higher proportion of eukaryote-eukaryote edges in Chinese network, which accounted for more than half of all connections (Fig. 2d). This dominance closely aligned with the observed depletion of core eukaryotic taxa and the proliferation of other lineages in Chinese lakes, suggesting that these opportunistic eukaryotes could rewired their associations to exploit newly available niches. Moreover, Chinese network exhibited higher transitivity, centralization, and assortativity, indicating a more complex and centralized architecture (Fig. 2e). By contrast, Tajikistani network had higher modularity and a greater proportion of negative links (Fig. 2d,e). Specifically, three of the four modules in Chinese network were dominated by eukaryotes (e.g., Chlorophyta and Cryptomycota), whereas three of the six modules in Tajik network were prokaryote-governed, largely by Proteobacteria (Supplementary Fig. 5). Because modularity can compartmentalize associations and negative links may dampen positive feedbacks, these properties are often interpreted as features that can buffer ecological perturbations in complex microbial systems^34,35^. The significant and opposite linear relationship between local clustering coefficient and degree between China and Tajikistan also indicated that highly connected hub taxa preferentially linked to one another to form a densely connected core in China (Fig. 2f), leading to a more pronounced rich-club effect (Fig. 2g), which could increase network integration but may also concentrate dependence on a small set of hub taxa^36^. These results suggested that Tajikistani network was simpler and more decentralized, which often represents its more stable characteristics^37^. This was further supported by greater network robustness in Tajikistan than in China, as evidenced by a shallower decline in natural connectivity under random node removal (Fig. 2h).

**Figure 2.**
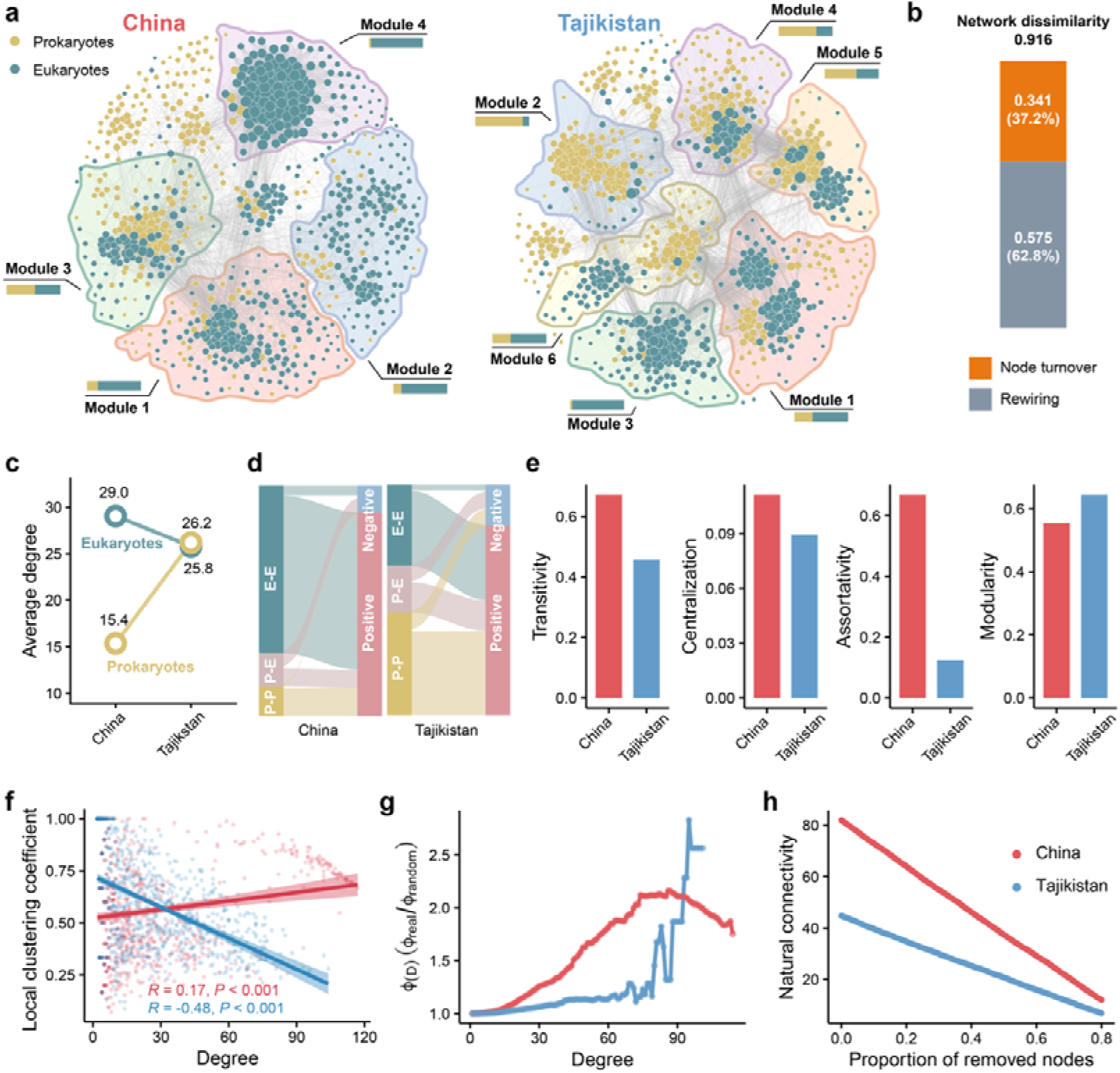
Regional differences of cross-kingdom microbial networks. **a**, Co-occurrence networks of microbial communities in Chinese and Tajikistani lakes. Shaded polygons encompass distinct modules. The accompanying stacked barplots denote the relative proportions of prokaryotes and eukaryotes within each module. **b**, Decomposition of the total network dissimilarity into rewiring and node turnover components. **c**, Comparison of the node average degree between eukaryotes and prokaryotes. **d**, Compositional shifts in edges, categorized by interaction types (E-E, Eukaryote-Eukaryote; P-E, Prokaryote-Eukaryote; P-P, Prokaryote-Prokaryote) and correlation signs (positive and negative). **e**, Topological properties, including transitivity, Centralization, assortativity, and modularity. **f**, Linear regressions between local clustering coefficient and node degree. **g**, Rich-club architecture. h, Network robustness, assessed by the decline of natural connectivity under random node removal.

We also focused on the differences in cross-kingdom links across regions. Comparative null-model analyses revealed that specific algae-bacteria linkages were highly region-dependent rather than phylogenetically conserved (Supplementary Fig. 6). For instance, negative links between Proteobacteria and some algal clades, including Phragmoplastophyta and Chlorophyta, were significantly enriched in Chinese lakes but depleted in Tajikistan. Such context dependency is plausible because associations between algae and bacteria could range from metabolite exchange and nutrient provisioning to competition, antagonism, and other negative interactions^38,39^. Furthermore, the bipartite networks of prokaryotes and eukaryotes showed greater connectance and nestedness in Tajikistan (Supplementary Fig. 7a). Compared to randomly removing intra-kingdoms links, removing cross-kingdom edges led to a faster decline in the natural connectivity of Tajikistani network but resulted in a slower decline in the natural connectivity of Chinese network (Supplementary Fig. 7b). This suggests a stronger dependence of Tajikistani network on cross-kingdom associations, which played an indispensable role in maintaining microbial community stability.

## Drivers of microbial co-occurrence patterns

We first explored which taxa were most prone to rewiring in networks and what constraints governed this process. The results showed that node rewiring exhibited significant phylogenetic signals in both kingdoms, with eukaryotes (λ = 0.72) exhibiting stronger phylogenetic constraints than prokaryotes (λ = 0.24) (Fig. 3a). This indicates that closely related taxa tended to exhibit similar rewiring behavior, especially among eukaryotic microorganisms. Interestingly, prokaryotic rewiring was predominantly driven by the recruitment of region-unique neighbors, while eukaryotic rewiring was mainly characterized by the reconfiguration of linkages among shared taxa (Supplementary Fig. 8). Such kingdom-specificity is consistent with the stronger conservation of many eukaryotic traits, including cell size, morphology, trophic mode, and physiological strategy^40–42^, whereas shorter generation times, higher passive dispersal potential, and greater functional plasticity may allow prokaryotes to more readily interact with locally recruited taxa under changing environmental conditions^43^. Furthermore, these reconnected patterns were often accompanied by significant changes in regional community attributes, indicating that the node rewiring was also closely related to the ecological generalization ability of the nodes and their positions within the networks (Fig. 3a). Random forest models identified maximum degree, salinity optimum, and niche width as the primary predictors of rewiring score (Fig. 3b), and all of them were significantly and negatively correlated with this score (Fig. 3c). Therefore, rewiring was not concentrated in highly connected ecological generalists, but instead tended to occur in less connected and more specialized taxa. Highly connected generalists usually occupy central network positions and provide a stable topological backbone for microbial communities^44^, making their association profiles less variable across regions. On the other hand, taxa adapted to lower-salinity conditions were especially sensitive to network reorganization, consistent with the strong role of salinity and ionic composition in structuring plateau lake microbiomes^13,17^.

**Figure 3.**
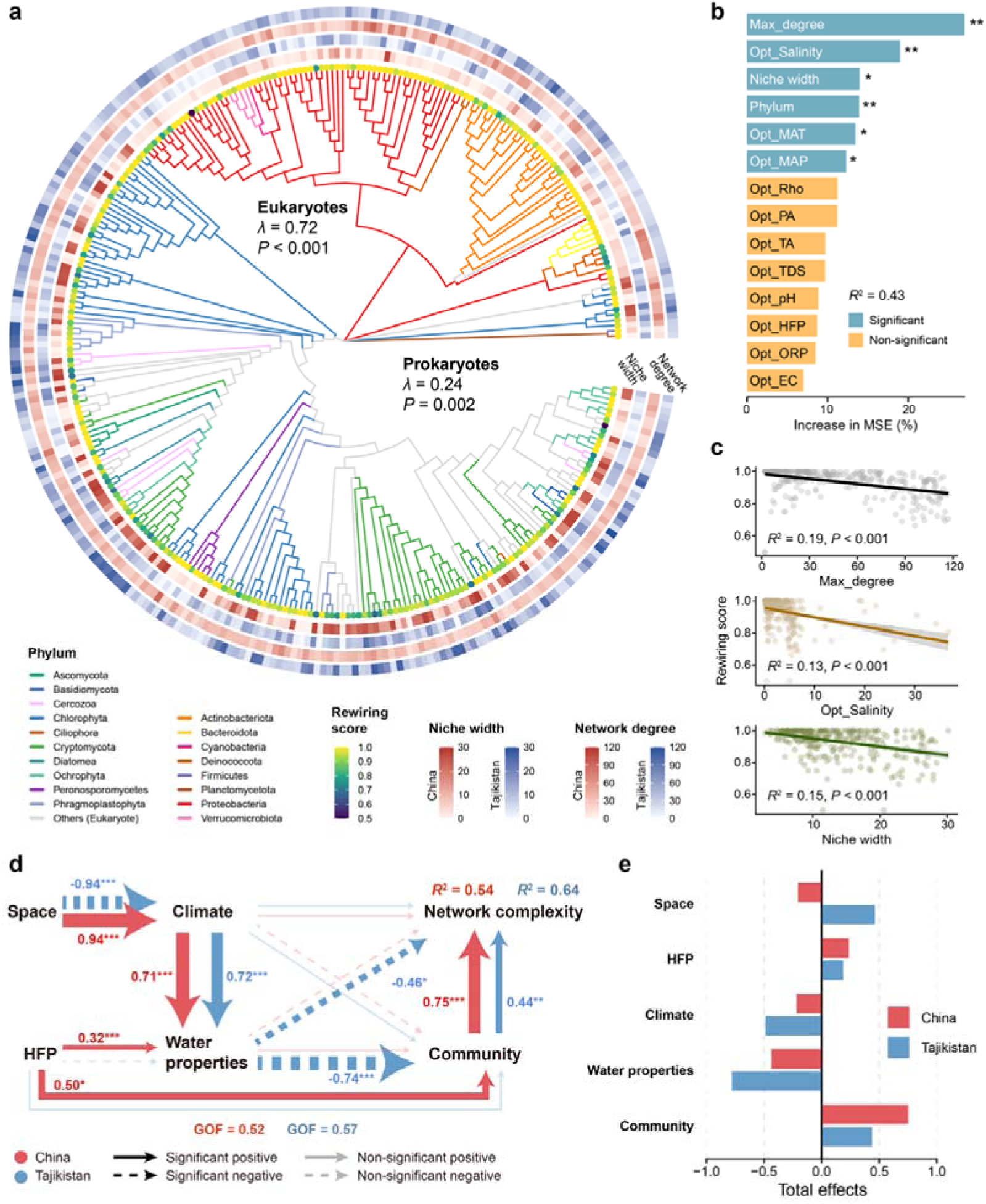
Drivers of microbial network divergence. a, Phylogenetic tree of shared microbial taxa. Pagel’s λ indicates the phylogenetic signal of node rewiring scores for eukaryotes and prokaryotes. Circles on the tree tips represent the rewiring scores. The surrounding concentric heatmaps display the rewiring score, niche width, and network degree of corresponding nodes in Chinese and Tajikistani lakes. **b**, Random forest models showing the importance of predictors on rewiring score. Predictor importance is measured as the percentage increase in mean squared error (MSE).

Asterisks indicate statistical significance (\*\*\**P* < 0.001, \*\**P* < 0.01, and \**P* < 0.05). **c**, Linear regression models between the rewiring score and maximum degree, optimal salinity, and niche width of each node. d, PLS-PM disentangling the effects of space, climate, HFP, water properties, and community dynamics on network complexity. Red and blue arrows represent pathways in Chinese and Tajikistani lakes, respectively. **e**, Total effects of each predictor pool on network complexity, derived from the PLS-PM analysis.

Because local rewiring may alter the functional compatibility of neighboring taxa, we further evaluated potential metabolic support around rewired prokaryotic nodes using genome-scale metabolic models reconstructed from matched reference genomes (See details in ***Methods***). We found metabolic complementarity in Tajikistan was significantly but slightly higher than that in China (Supplementary Fig. 9a). Ordination of metabolite exchange profiles showed that country explained only a very small proportion of the variation (*R*^2^ = 0.005), whereas phylum pairing explained substantially more (*R*^2^ = 0.076) (Supplementary Fig. 9b). This indicated that the metabolic support architecture of rewired nodes was largely preserved, but it was also constrained by the region. For instance, the potential transfer of magnesium, D-glyceraldehyde, and D-glutamate was significantly more frequent in Chinese lakes, while Tajikistani networks were characterized by a significantly higher frequency of transfer for L-lactaldehyde, potassium, and sulfite (Supplementary Fig. 9c). Altogether, these findings suggested that node rewiring was likely driven by the necessity to maintain or recalibrate metabolic complementarity under regional environmental stress^45,46^.

We then examined how regional environmental forces shaped network complexity using partial least squares path modelling (PLS-PM). In Tajikistani lakes, climate influenced water properties, which in turn exerted a significant direct constraint on network complexity (Fig. 3d). Water properties therefore emerged as the dominant driver of network complexity (Fig. 3e), reflecting an environment-controlled ecological regime. This pattern is consistent with previous plateau lake studies showing that physicochemical properties of water (e.g., salinity and conductivity) act as major filters of microbial community and co-occurrence structure^17,47^. In Chinese lakes, by contrast, the direct link between water properties and network complexity was no longer detectable. Human footprint directly affected both water properties and microbial communities (Fig. 3d), and community dynamics became the strongest predictor of network complexity (Fig. 3e). These observations indicated that anthropogenic influence did not simply act through water conditions, but also through reshaping the microbial species pool and cross-kingdom associations, because human activities could alter dissolved organic matter content and water quality and introduce distinct bacterial signatures to restructure microbial co-occurrence networks^48–50^. The Spearman’s correlation analysis confirmed our findings that the environmental variables and network complexity were closely linked in Tajikistan but decoupled in China (Supplementary Fig. 10a). The random forest model also showed that cross-kingdom edges and microbial diversity were the strongest predictors of network complexity in China. In contrast, the complexity variation of Tajikistani network was more explained by environmental variables, such as total dissolved solids (TDS), electrical conductivity (EC), and salinity (Supplementary Fig. 10b). Notably, temperature anomalies (TA) and precipitation anomalies (PA) significantly influenced network complexity exclusively in Tajikistan, indicating that microbial association patterns in this less disturbed region may still retain climatic legacies. In Chinese lakes, stronger human footprint appeared to override these historical environmental signals. As expected, mantel correlations between environmental distance and network topology distance revealed a pronounced collapse driven by the human footprint (HFP) (Supplementary Fig. 10c). Energy landscape modeling further elucidated that increased HFP reconfigured the microbial assembly energy landscape by destabilizing the originally dominant low-energy state (Supplementary Fig. 10d), indicating a reduction in community stability^51^. These results emphasized that human activities played a crucial role in affecting microbial networks.

### Microbial element cycling functions and potential biological safety risks

PCoA revealed significant geographic divergence in microbial functional potentials across all four element cycles, including carbon, nitrogen, phosphorus, and sulfur cycling (Fig. 4a). Systemic functional profiling demonstrated that biogeochemical pathways were predominantly enriched in Chinese lakes (Fig. 4b). For carbon cycling, pathways related to carbon fixation and release, as well as organic degradation and biosynthesis were significantly more abundant in China, suggesting stronger potential for both biomass production and organic matter processing. Regarding nitrogen cycling, denitrification, anammox, assimilatory nitrate reduction, and nitrification, were highly enriched in China. These pathways can accelerate reactive nitrogen turnover and may increase the potential for nitrogen loss and nitrogen-related greenhouse gas production under redox-variable conditions^52^. Purine, pyrimidine, and pyruvate metabolism, alongside transporters, two-component systems, organic phosphoester hydrolysis, and phosphotransferase system were also enriched in China, together with broad sulfur disproportionation, sulfur oxidation, and dissimilatory sulfur reduction and oxidation. These patterns suggested that Chinese lake microbiomes possessed greater metabolic versatility, exhibiting an expanded capacity to couple carbon turnover with nitrogen and sulfur redox transformations^13,53^. This broad functional enrichment was likely driven by the stronger human footprint and allochthonous hydrological inputs in Chinese lakes, which could alter dissolved organic matter profiles, enhance nutrient availability, and increase redox heterogeneity, thereby creating more opportunities for microorganisms involved in element cycling^54,55^. Instead of broad enrichment of transformation pathways, lake microbiomes in Tajikistan were characterized by higher nitrogen fixation and stronger phosphonate and phosphinate metabolism (Fig. 4b). This observation is consistent with nutrient-limited systems, where microorganisms invest in pathways that supplement bioavailable nitrogen and access alternative phosphorus sources^56,57^.

**Figure 4.**
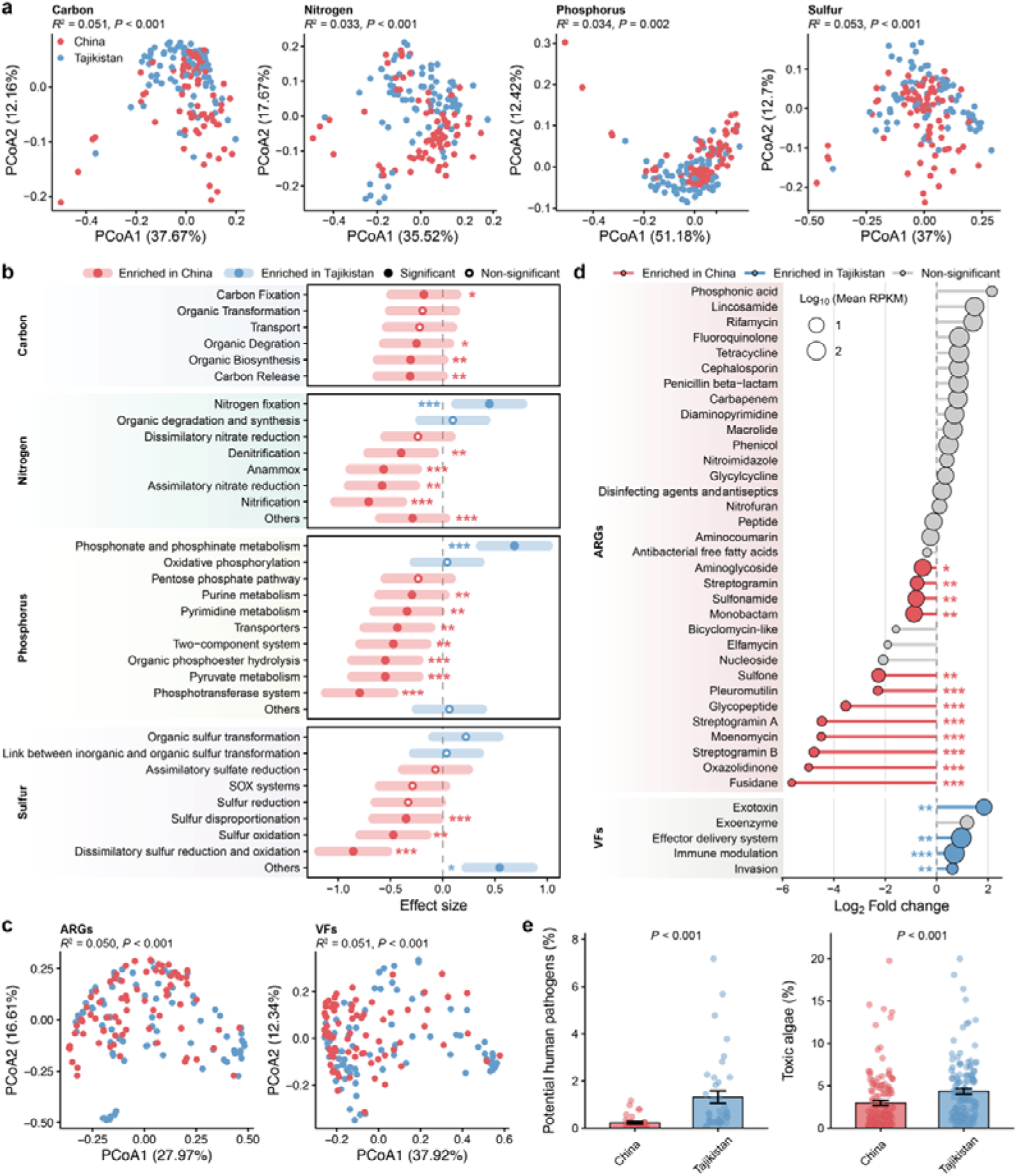
Microbial element cycling functions and potential biosafety risks. **a**, PCoA with PERMANOVA illustrating the structural divergence of functional genes associated with carbon, nitrogen, phosphorus, and sulfur cycling between Chinese and Tajikistani lakes. **b**, Differential enrichment of specific biogeochemical pathways. Error bars represent the mean ± one standard deviation. **c**, PCoA with PERMANOVA for ARGs and VFs. **d**, Differential abundance of specific ARG classes and VF categories. **e**, Relative abundance of potential human pathogens and toxic algae. Bar heights represent the mean values, and error bars indicate the standard error. *P* values were calculated using two-sided Wilcoxon rank-sum tests. Asterisks indicate statistical significance (\*\*\**P* < 0.001, \*\**P* < 0.01, and \**P* < 0.05).

Differential abundance analysis at the gene level corroborated these shifts (Supplementary Fig. 11). We found 974 genes were significantly enriched in China (e.g., *cbiF*, *bamB*, and *pdhC*) compared to 211 in Tajikistan (e.g., *bchE*, *thrH*, and *vhtG*). For nitrogen cycling, key genes driving ammonia/methane oxidation and nitrification (e.g., *amoA*_A, *amoC_A*, *hao*, and *pmoA*) were conspicuously enriched in China, whereas the nitrite reduction gene *nrfC* was favored in Tajikistan. Genes of phosphorus cycling such as *aepW*, *pphA*, and *purP* were significantly enriched in China, while a suite of genes governing phosphonate metabolism (e.g., *phnM*, *phpC*, *phnT*, *fomC*) were abundant in Tajikistan. A total of 61 sulfur cycling genes were differentially enriched, with 49 genes enriched in China and 12 in Tajikistan. Among them, specific sulfur utilization and degradation genes like *metA*, *dmdA*, and *soeB* exhibited high enrichment in Tajikistani lakes.

To rigorously assess potential biosafety risks, we applied the antibiotic resistance gene (ARG) risk framework established by Zhang et al., which comprehensively integrates human accessibility, mobility, pathogenicity, and clinical availability^58^. Based on it, we identified 1,132 risky ARGs. Meanwhile, we selected five classes of potentially offensive virulence factors (VFs) for subsequent analysis. PCoA demonstrated significant differences for both ARG and VF profiles between China and Tajikistan (Fig. 4c). Chinese lakes harbored a profoundly expanded risky ARG repertoire, featuring significant enrichments in resistance to fusidane, oxazolidinone, streptogramin A/B, moenomycin, glycopeptide, pleuromutilin, and sulfone (Fig. 4d). Conversely, Tajikistani lakes, despite possessing contracted risky ARGs, exhibited a marked enrichment in core offensive VFs, including exotoxin, invasion, immune modulation, and effector delivery system (Fig. 4d). Consistent with the VF profiles, Tajikistani lakes hosted a significantly higher relative proportion of both potential human pathogens and toxic algae (Fig. 4e). Together, these observations showed that Chinese lakes were characterized mainly by resistance-related risks. The expanded risky ARG repertoire in Chinese region was likely associated with stronger human footprint, which could introduce antibiotics, resistant bacteria, mobile genetic elements, and clinically relevant ARGs into lakes through wastewater, runoff, tourism, livestock, and settlement-related inputs^18^. In contrast, Tajikistani lakes showed stronger pathogenicity- and toxicity-related potential, which could be attributed to the natural enrichment of taxa with hazards. Because environmental bacteria could maintain virulence-related traits, these traits may help them interact with eukaryotic hosts, avoid predation by microbial eukaryotes, or survive within host cells, even in the absence of strong human contamination^59,60^.

## Influence of microbial networks on potential ecological consequences

We normalized abundances of functional genes related to carbon, nitrogen, phosphorus and sulfur cycling and integrated them into microbial multifunctionality (MMF). Similarly, the normalized abundances of ARGs, VFs, potential human pathogens, and toxic algae were combined into potential biosafety risk (PBR). The results showed that Chinese lakes exhibited significantly higher MMF but lower PBR compared to Tajikistani lakes (Fig. 5a), indicating that microbial functional enhancement and biosafety risk were not necessarily coupled in the same direction. Mantel tests confirmed that the variations of MMF and PBR were correlated not only with taxonomic turnover but also with network dissimilarity, highlighting the indispensable role of rewiring-dominated microbial network in regulating these ecological consequences (Fig. 5b). Furthermore, rewiring score of each node was positively associated with its effect size on MMF (*P* = 0.055) but negatively correlated with the effect size on PBR (*P* < 0.001) (Fig. 5c), suggesting that highly rewired nodes tended to contribute more to ecosystem functioning while being less associated with biosafety risk. This reflects that network rewiring could mediate the community-level trade-off between MMF and PBR. To verify this discovery, we first trained a graph embedding-random forest framework that achieved moderate predictive performance for MMF and PBR (*R*^2^ = 0.27 and 0.43, respectively) (Supplementary Fig. 12a). We then performed an *in silico* rewiring analysis, in which local node neighborhoods were progressively rewritten toward the opposite regional template (See details in ***Methods***). Two representative Tajikistani subnetworks with low MMF and high PBR, as well as two Chinese subnetworks characterized by high MMF and low PBR were selected for simulations. As expected, rewiring Tajikistani subnetworks toward Chinese templates increased MMF and reduced PBR, whereas rewiring Chinese subnetworks toward Tajikistani templates produced the opposite shifts (Supplementary Fig. 12b). Consequently, the function-risk trade-off reflects a spatial segregation of distinct ecological configurations. The network rewiring we observed acts as the primary mechanistic vehicle translating these regional selective pressures into divergent potential ecological outcomes, thereby giving rise to a macro-scale mismatch between potential ecosystem service and biological safety burden.

**Figure 5.**
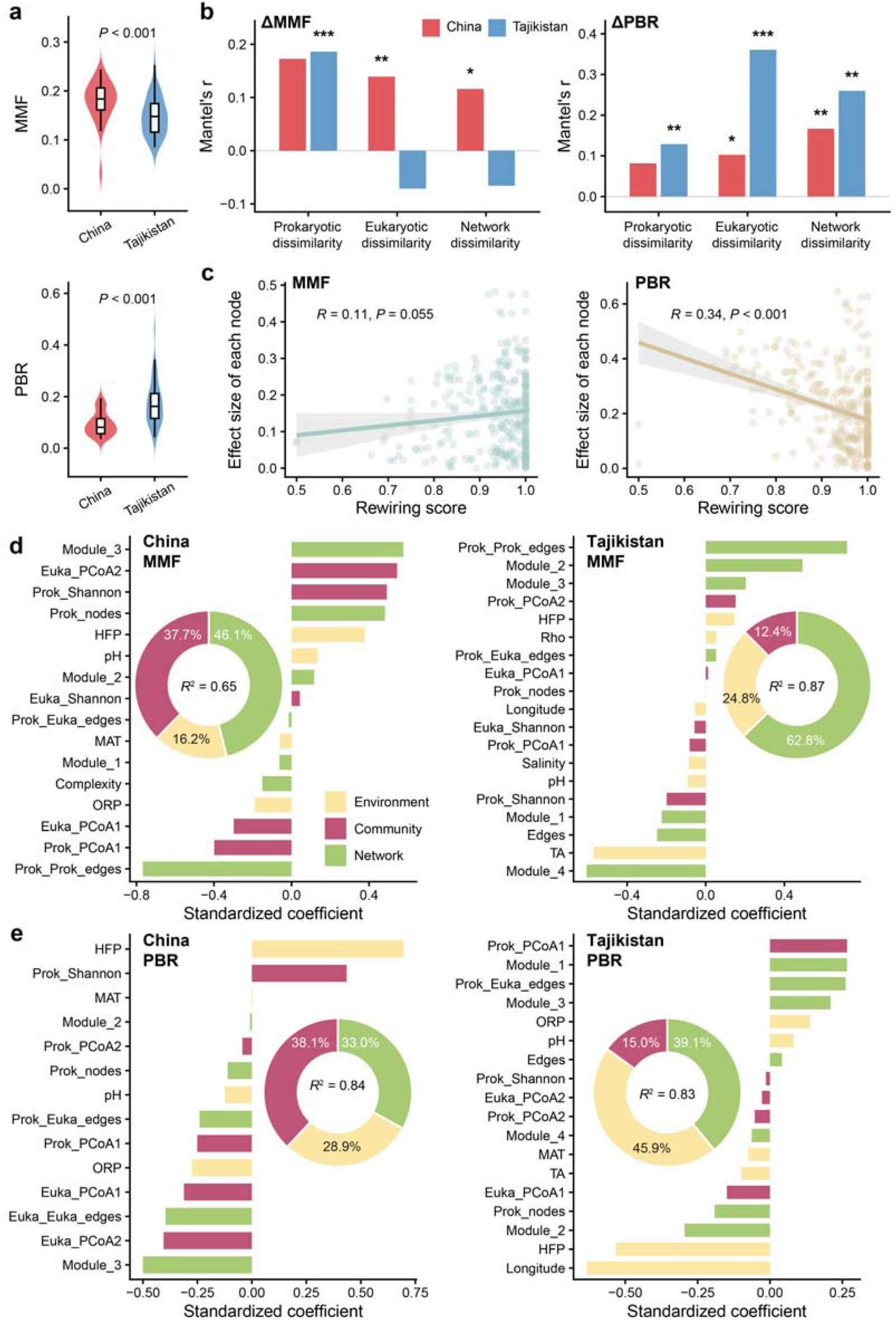
The role of microbial networks in mediating MMF and PBR. **a**, Violin plots comparing the MMF and PBR between Chinese and Tajikistani lakes. Hinges show the 25th, 50th, and 75th percentiles. *P* values were calculated using two-sided Wilcoxon rank-sum tests. **b**, Mantel correlations (*r*) evaluating the associations between the variations of ecological consequences (ΔMMF and ΔPBR) and the prokaryotic, eukaryotic, and network dissimilarities. Asterisks indicate statistical significance (\*\*\**P* < 0.001, \*\**P* < 0.01, and \**P* < 0.05). **c**, Linear regression models assessing the relationship between the rewiring score of individual nodes and their corresponding effect sizes on MMF and PBR. Complete GLMs identifying the drivers of (**d**) MMF and (**e**) PBR. The inset donut charts display the total variance explained by the complete models, with segments quantifying the relative percentage contributions of the three predictor pools to the explained variance.

To determine whether network properties improved the explanation of MMF and PBR beyond environmental and community variables, we compared two generalized linear model (GLM) frameworks after removing collinear predictors within each region (Supplementary Fig. 13). The first model included only environmental and community variables, whereas the second additionally included network properties. Incorporating network properties consistently improved the explanation of variance in both MMF and PBR and substantially altered the contribution structure. For MMF, the variance explained increased from *R*^2^ = 0.56 to 0.65 in China and from 0.69 to 0.87 in Tajikistan, with network variables becoming the largest contributor in both regions, particularly in Tajikistan (Fig. 5d and Supplementary Fig. 14a). This supports the view that microbial diversity and network complexity could jointly predict ecosystem functioning, because ecosystem processes often depend on complementary taxa and coordinated functional guilds rather than isolated community members^61,62^. In terms of PBR, the explanation increased from *R*^2^ = 0.75 to 0.84 in China, where community and network effects were of comparable importance. Tajikistan model remained dominated by environment and network variables (*R*^2^ = 0.83) and showed only a minor contribution from community attributes (Fig. 5e and Supplementary Fig. 14b).

Notably, several predictors exhibited clear region-specific regulatory effects. Specifically, prokaryote-prokaryote edges showed opposite effects between regions for MMF, being strongly negative in China but positive in Tajikistan (Fig. 5d), indicating that similar topological features might support multifunctionality through different ways. In Chinese lakes, dense prokaryote-only associations may reflect redundancy, competition, or stress-linked clustering under stronger anthropogenic pressure, whereas in Tajikistani lakes they may coordinate nutrient acquisition and element cycling under oligotrophic conditions. Likewise, for PBR, cross-kingdom edges were negatively associated with PBR in China but positively associated with risk in Tajikistan (Fig. 5e), suggesting that prokaryote-eukaryote interactions could either buffer or amplify biosafety risk under distinct conditions. HFP also exhibited strong context dependence for PBR, showing the greatest positive effect in China but a strong negative impact in Tajikistan. This contrast suggests that biosafety responses to human pressure cannot be generalized across the Pamir Plateau, because the same pressure may interact with different microbial species pools, network structures, and background environmental conditions. Moreover, network modules contributed significantly to these ecological outcomes, for example, Module_3 markedly increased MMF but decreased PBR in China. This implies that microorganisms could leverage highly cohesive functional modules to synergistically regulate ecosystem services^63^. Overall, our study highlights the importance of incorporating microbial interaction networks into high-altitude lake assessment, because ecosystem functioning and biosafety risk depend not only on which microorganisms are present, but also on how they are connected.

To our knowledge, this study provides the first cross-border assessments of lake prokaryotic and eukaryotic microbiomes on the Pamir Plateau and advances our understanding of high-altitude microbial biogeography beyond conventional compositional shifts to the underlying network rewiring. More importantly, our results identify network rewiring as a pathway that translates regional environmental divergence into divergent potential ecological outcomes by mediating the trade-off between microbial multifunctionality and potential biosafety risk. However, several limitations should be acknowledged. First, our sampling design could not fully capture spatial heterogeneity across the entire lake surface, shoreline-offshore gradients, or vertical water-column structure. In addition, samples were collected within a limited sampling window, and therefore seasonal succession, interannual variability, and episodic hydrological events remain unresolved. Second, because cross-border sampling in remote high-altitude mountain regions is logistically challenging^11^, we were unable to measure a broader suite of physicochemical and process-level variables. Measurements of greenhouse gas emissions, variables related to carbon, nitrogen, phosphorus, and sulfur would be valuable for validating the functional implications inferred from metagenomic profiles. Future work combining spatially stratified sampling, long-term time-series monitoring, expanded environmental measurements, and direct process-rate assays will further clarify how microbial biogeography translates into ecosystem functioning and water biosafety in high-altitude lakes.

## Conclusion

Our study demonstrates that regional divergence across the Pamir Plateau reshapes high-altitude lake microbiomes by altering prokaryotic and eukaryotic biogeography and rewiring microbial associations, thereby triggering distinct potential ecological consequences (Fig. 6). Tajikistani lakes maintain higher prokaryotic diversity and stronger eukaryotic connectivity, whereas Chinese lakes show higher eukaryotic diversity and greater eukaryotic dispersal limitation. These divergent biogeographic patterns lead to distinct network characteristics through interaction rewiring. Tajikistani networks are more modular and stable under strong environmental filtering, while Chinese networks become more centralized and complex, with human footprint weakening the coupling between environmental conditions and network structure. Functionally, Chinese lake microbiome exhibited higher microbial multifunctionality, while Tajikistani lakes showed stronger potential biosafety risk, suggesting a trade-off between them mediated by network rewiring. Taken together, our study provides new insights into the microbial responses of high-altitude lake microbiomes to regional environmental divergence and motivates network-informed monitoring of transboundary freshwater systems under global change.

**Figure 6.**
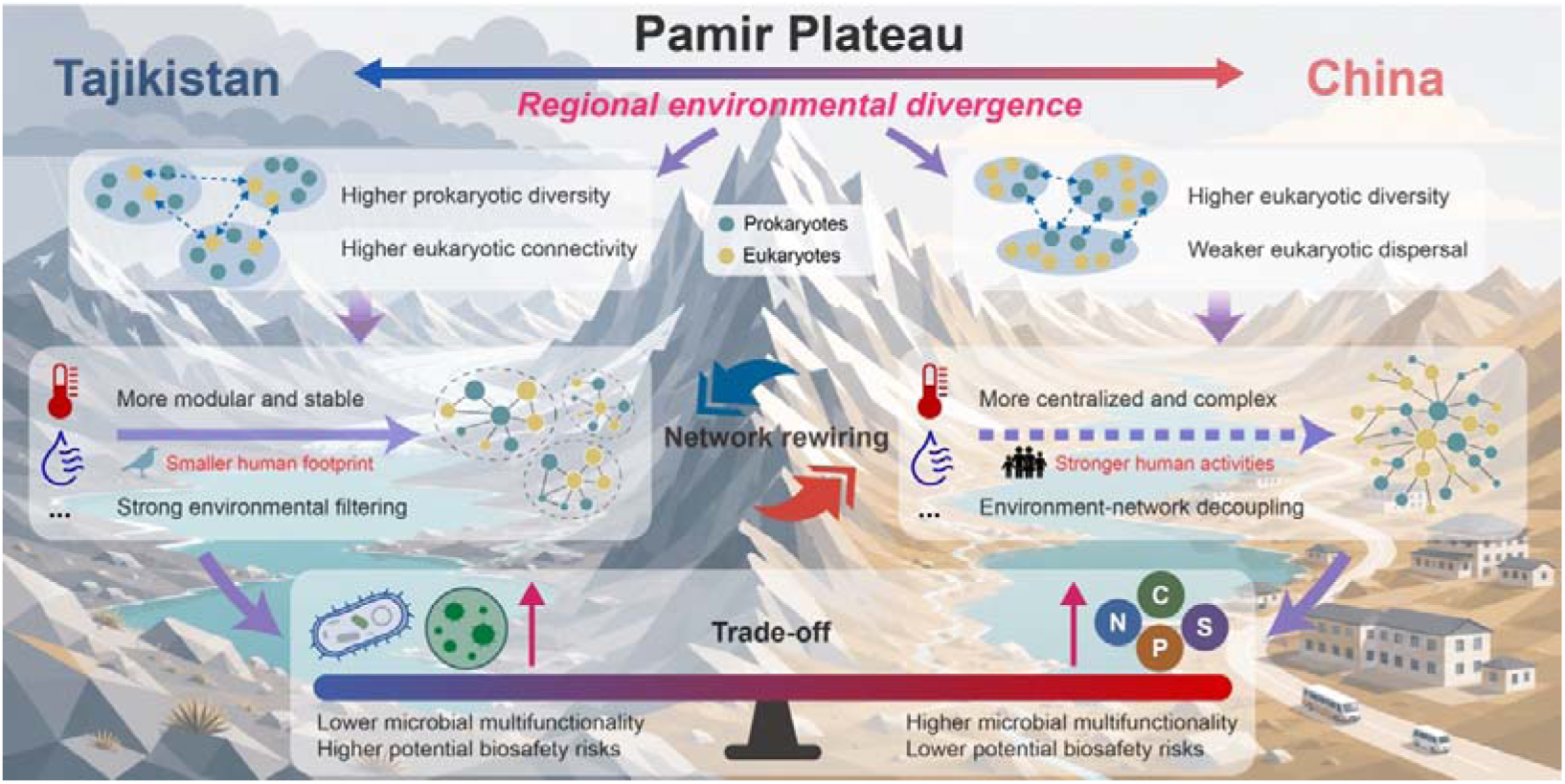
Conceptual model linking microbial biogeography to ecological consequences by network rewiring. Regional environmental divergence across the Pamir Plateau generated contrasting microbial biogeographic patterns between Tajikistani and Chinese lakes. Tajikistani lakes were characterized by higher prokaryotic diversity and stronger eukaryotic connectivity, whereas Chinese lakes showed higher eukaryotic diversity but weaker eukaryotic dispersal. These divergent biogeographic responses led to distinct cross-kingdom network architectures. In Tajikistani lakes, stronger environmental filtering was linked to more modular and stable microbial networks. In Chinese lakes, stronger human activities weakened the coupling between environmental conditions and network structure, leading to more centralized and complex networks. Network rewiring acted as a bridge between regional biogeography and ecological consequences, mediating a trade-off between microbial multifunctionality and potential biosafety risks across the Pamir Plateau.

## Methods

### Sampling sites and collection

Field sampling was conducted in July and August 2025 in 20 high-altitude lakes across the Pamir Plateau, spanning both Chinese (n = 10) and Tajikistani (n = 10) regions. These sampling lakes ranged from 37.45°N to 39.42°N and from 72.16°E to 75.58°E. Detailed geographic information can be found in Supplementary Table 1. At each lake, surface water samples were collected approximately 20 m from the lake margin at a depth of 10-30 cm. Ten replicate samples were collected from each lake, with adjacent sampling points separated by approximately 100 m. For each replicate, 2 L of lake water was collected using sterile water-sampling bags.

Water samples were processed within several hours after collection. Each sample was first passed through a 200 µm filter to remove insoluble impurities, large organisms, aquatic plants, and other debris. The filtrate was then sequentially filtered through 3 µm and 0.2 µm polycarbonate membranes (Millipore, Billerica, MA, USA) to retain size-fractionated microbes for downstream DNA extraction and sequencing.

### Water properties

Water physicochemical properties, including pH, oxidation-reduction potential (ORP), electrical conductivity (EC), electrical resistivity (Rho), total dissolved solids (TDS), and salinity, were measured *in situ* at each sampling point. These measurements were recorded simultaneously with sample collection using a portable multiparameter water quality analyzer (HI9828 HANNA, Italy), which was calibrated according to the manufacturer’s specifications prior to deployment.

### Large-scale environmental and paleoclimatic drivers

Climatic variables were extracted based on the precise geographical coordinates of each sampling point. Mean annual temperature (MAT) and mean annual precipitation (MAP) were retrieved from the WorldClim database (https://www.worldclim.org/) at a spatial resolution of 30 arc seconds (∼1 km). Furthermore, given the potential legacy effects of paleoclimate in shaping modern microbial diversity patterns, historical MAT and MAP during the last glacial maximum (LGM, ∼22 ka BP) were acquired from the CHELSA project^64^. CHELSA-TraCE21k data provides climatic variables at 30Oarc seconds spatial resolution in 100-year time steps for the last 21,000Oyears. To quantitatively represent the magnitude of past climate change, TA and PA for each sampling point were calculated as the absolute differences in MAT and MAP, respectively, between the LGM and contemporary periods. HFP scores were extracted from the global Human Footprint dataset with 1 km2 resolution^65^. This dataset integrates eight anthropogenic pressure layers, including built environments, population density, night-time lights/electric infrastructure, croplands, pastures, roads, railways, and navigable waterways.

### DNA extraction

Total DNA for amplicon and metagenomic sequencing was extracted from water sample filters using the PF Mag-Bind Stool DNA Kit (Omega Bio-Tek, USA) and the MolPure Mag96 Soil/Stool DNA Kit (Yeasen, China), respectively, following the manufacturers’ instructions. After extraction, DNA concentration and purity were assessed using a NanoDrop 2000 spectrophotometer and a Fluo-800 fluorometer. DNA integrity was evaluated using 1% agarose gel electrophoresis.

### Amplicon sequencing

The prokaryotic and eukaryotic microbial communities were profiled by amplifying the 16S rRNA gene and 18S rRNA gene, respectively. The V3-V4 region of the 16S rRNA gene was amplified using the primer pair 338F (5’-ACTCCTACGGGAGGCAGCAG-3’) and 806R (5’-GGACTACHVGGGTWTCTAAT-3’), whereas the V4 region of the 18S rRNA gene was amplified using V4F (5’-CGGTAATTCCAGCTCCAATA-3’) and V4R (5’-GATCCCCHWACTTTCGTTCTTGA-3’). Triplicate PCR products from each sample were pooled and checked by 2% agarose gel electrophoresis. Amplicons of the expected size were purified using a magnetic bead-based gel recovery kit and quantified using a Quantus fluorometer. Purified amplicons were pooled in equimolar amounts, and sequencing libraries were prepared using the NEXTFLEX Rapid DNA-Seq Kit (Bioo Scientific, USA). Paired-end sequencing was performed on an Illumina MiSeq PE300 platform.

Bioinformatics processing, including filtering, dereplication, sample inference, chimera identification, and merging of paired-end reads, was performed using the Divisive Amplicon Denoising Algorithm 2 (DADA2)^66^. The taxonomical annotation of the representative sequences of ASVs was performed with a naïve Bayesian classifier using the Silva v.138.1 database^67^. Samples were rarefied to the same sequence depth (30,000 reads for both prokaryotes and eukaryotes per sample). This resulted in some samples with insufficient reads being excluded, and the information on the remaining samples included in the subsequent analysis can be found in Supplementary Table 2.

### Metagenomic sequencing and quality control

High-quality DNA extracts were used for paired-end library construction and sequencing. Some of the samples were excluded because of their poor quality and the information on the remaining samples included in the subsequent analysis can be found in Supplementary Table 2. Qualified libraries were sequenced on the BGI DNBSEQ-T7 platform using a PE150 strategy. The average raw read output for each DNA sample was approximately 20 Gb. Raw paired-end reads were quality-filtered using Trimmomatic (v.0.39)^68^. Briefly, sequencing adapters were removed, low-quality bases at the 5’ and 3’ ends were trimmed, and reads were further scanned using a 5-bp sliding window, with trimming initiated when the average quality score within the window fell below 20. Reads shorter than 50 bp after trimming were discarded. Sequencing quality was further evaluated based on Q20, Q30, and read retention statistics. The retained high-quality paired reads were used for downstream metagenomic functional annotation.

### Functional gene annotation based on reads

To profile microbial biogeochemical potentials, we first established a carbon, nitrogen, phosphorus, and sulfur cycling database (CNPS cycling gene database) by integrating CCycDB (https://ccycdb.github.io), NCycDB^69^, PCycDB^70^, and SCycDB^71^. This database was rigorously dereplicated using a custom script based on sequence and gene name. For potential biosafety risk assessments, we downloaded the CARD^72^ and VFDB^73^ databases. Metagenomic sequencing reads were subsequently aligned against the custom CNPS cycling gene database, CARD, and VFDB utilizing the DIAMOND blastx algorithm (v 2.1.10), with the parameters “--evalue 1e-10, --id 80, --query-cover 80 and --sensitive, with --max-target-seqs 1” applied to eliminate redundant alignments^74^. Finally, the abundances of all successfully assigned functional genes, ARGs, and VFs were mathematically normalized and expressed as reads per kilobase per million mapped reads (RPKM). To focus on ARGs with greater potential ecological and health relevance, ARGs were filtered utilizing the risk evaluation framework proposed by Zhang et al.,^58^ successfully identifying 1,132 risky ARGs. Concurrently, five categories of potentially offensive VFs were chosen for downstream analysis.

### Assessment of potential human pathogens and toxic algae

The potential human pathogens were predicted based on the functional annotation of 16S rRNA gene sequences using the FAPROTAX database^75^. Specifically, the relative abundances of ASVs assigned to the functional categories “human_pathogens_all” and “human_pathogens_pneumonia” were aggregated to represent the total proportion of potential human pathogens in each sample. For the eukaryotic communities, the identification of toxic algae was conducted by querying the representative 18S rRNA sequences against the Harmful and Toxic Microalgae Database (HTMaDB; https://github.com/lisse0321/HTMaDB) using BLASTn. To ensure taxonomic accuracy, the alignments were strictly filtered using a minimum sequence identity threshold of 99% and an e-value threshold of < 1e-5. The relative abundances of the successfully matched eukaryotic ASVs were then summed to determine the total proportion of toxic algae in each sample.

## Statistical analyses

### Microbial community attributes

Prokaryotic and eukaryotic alpha diversity, including richness, Shannon, and Simpson indices, were calculated using the vegan R package^76^. PCoA based on Bray-Curtis dissimilarity was employed to visualize beta diversity, with PERMANOVA being applied to assess the effects using the adonis2 function in the vegan R package^76^. The ALDEx2 R package was used to determine differences at the ASV level in terms of relative abundance^77^. For taxa passing filtration (mean relative abundance > 0.01% and occurrence rate > 10%), we applied the aldex function with default settings (mc.samples = 128, test = “t”, denom = “all”). Wilcoxon rank-sum tests followed by Benjamini–Hochberg false discovery rate (FDR) correction was then implemented via the aldex.ttest function to identify ASVs showing significantly different relative abundances between China and Tajikistan. Levins’ niche width was calculated using the spaa R package^78^. Microbial communities were partitioned into core and peripheral (specific, unique, and other) sub-communities. Specifically, core taxa were defined as ASVs occurring in > 50% with relative abundance > 0.1% of all samples; specific taxa were those restricted to a single lake but detected in multiple samples; unique taxa were ASVs exclusively found in a single sample; and all remaining ASVs were categorized as other taxa. DDR were assessed by calculating the linear regression between geographic distance and community dissimilarity. Procrustes analysis was conducted to evaluate the associations between prokaryotic and eukaryotic communities using the procrustes function in the vegan R package^76^. To determine microbial connectivity between lakes, source tracking was performed using the FEAST R package^79^. A null-model-based framework was utilized to quantify the relative contributions of deterministic and stochastic processes governing community assembly using the iCAMP R package^80^.

### Microbial co-occurrence networks

Cross-kingdom co-occurrence networks were constructed separately for Chinese and Tajikistani lakes using the SparCC algorithm by the SpiecEasi R package^81^. For 18S rRNA gene datasets, ASV tables from the 3 µm and 0.2 µm fractions were merged by summing ASV abundances within each lake before constructing networks. The SparCC results were filtered by the thresholds |*r*| > 0.5 and *P* < 0.05. To quantify the divergence between Chinese and Tajikistani networks, we performed an edge-level decomposition of network dissimilarity. We used the broad network rewiring framework of Ward et al. in which rewiring refers to topological reorganization of ecological networks caused by node gain or loss and/or changes in link patterning^24^. For each regional network, edges were represented as unordered node pairs. Total network dissimilarity was calculated as the proportion of non-shared edges between the two networks, using β = (A + B - 2J)/(A + B), where A and B are the numbers of edges in the Chinese and Tajikistani networks, respectively, and J is the number of edges shared by both networks. Nodes were classified as shared if they occurred in both regional networks, or region-specific if they occurred in only one network. Rewiring was defined as non-shared edges involving at least one shared node, including both changes in associations among shared taxa and the recruitment of region-specific neighbors by shared taxa. Node turnover was defined as non-shared edges connecting two region-specific nodes. The relative contribution of each component was calculated by dividing its edge-dissimilarity component by total network dissimilarity. For each node shared by the Chinese and Tajikistani networks, we further calculated a rewiring score to quantify the extent to which its association neighborhood changed between regions. The score was defined as the Jaccard distance between the node’s neighbor set in the Chinese network and its neighbor set in the Tajikistani network. The phylogenetic signal of the rewiring scores was quantified using Pagel’s λ statistic using the phytools R package^82^. Network properties including transitivity, modularity, degree, centralization, and assortativity were calculated using the igraph R package^83^. Bipartite networks between prokaryotes and eukaryotes were extracted to calculate connectance and nestedness. The rich-club effect was evaluated to ascertain whether highly connected hub taxa preferentially interact with one another to form a densely interconnected structural core^36^. We computed the rich-club coefficient, defined as the ratio of observed edges among nodes with a degree greater than *k* to the maximum possible number of edges among them. To control for structural artifacts inherently caused by high-degree nodes, this empirical coefficient was normalized against the average coefficient derived from 999 randomized networks with identical degree distributions. A normalized coefficient significantly greater than 1 across the topological gradient confirms a pronounced rich-club architecture within the network. To determine whether specific cross-kingdom interactions, particularly linkages between prokaryotic and algal phyla, were deterministically enriched or depleted, we employed a comparative null-model framework. For each regional interactome, we generated 999 randomized networks using a degree-preserving edge-swapping algorithm. The observed number of cross-kingdom edges between specific clades was then compared against the null distribution to calculate standard effect sizes (Z-scores). Network robustness was assessed by simulating the decline in natural connectivity under targeted and random node/edge removal (n = 100). The statistical significance of the phylogenetic signal was assessed through a likelihood ratio test, which compared the fitted model against a null model assuming no phylogenetic signal. Energy landscape modeling was applied to evaluate changes in microbial stability along the gradient of HFP using the rELA R package^51^.

## Metabolic complementarity

To elucidate the metabolic mechanisms underlying network rewiring, we reconstructed GEMs for the prokaryotic network members. Representative genomes corresponding to the prokaryotic 16S rRNA ASVs were retrieved from the NCBI and GTDB reference databases, applying a stringent sequence identity threshold of >99%. GEMs were generated using CarveMe under default parameters with a gap-filling procedure performed on media lysogeny broth (LB)^84^. To assess the potential for metabolic cross-feeding between interacting taxa, we calculated the metabolic complementarity index using PhyloMint^85^. This index quantifies the theoretical capacity of a recipient strain (Strain A) to utilize the metabolic byproducts secreted by a donor strain (Strain B). Potentially transferable metabolites were defined as the intersection between the seed set of Strain A and the non-seed set of Strain B. According to this framework, we first compared the profile difference of potentially transferable metabolites across regions using PCoA based on Jaccard dissimilarity, with PERMANOVA being employed to assess the effects using the adonis2 function in the vegan R package^76^. We also quantified the specific metabolite exchange frequencies between China and Tajikistan. Fisher’s exact test was employed to determine whether the transfer of specific metabolites was significantly enriched or depleted in a region-specific manner. For each identified exchangeable metabolite, a 2 x 2 contingency table was constructed based on its presence or absence across all directed cross-feeding edges within the respective regional networks. To control for false positives arising from multiple testing, raw P values were subsequently adjusted using the Benjamini-Hochberg FDR procedure.

### Partial least squares path modeling

Using the plspm R package^86^, partial least squares path modeling was applied to explore the direct and indirect effects of space, human activities, climate, water properties, and microbial community on network complexity. Network complexity was obtained by normalizing transitivity, modularity (its reciprocal), degree, centralization, and assortativity and then averaging them. Notably, we defined box “Community” as a comprehensive variable integrating both taxonomic diversities (prokaryotic and eukaryotic Shannon indices) and the number of cross-kingdom edges to represent the dynamics of microbial communities mixing. To ensure the robustness of the model, we utilized the Goodness-of-Fit (GoF) index to evaluate the overall predictive power.

### Mantel test

Mantel tests were employed at two distinct levels to evaluate the associations between different ecological distance matrices using the vegan R package. First, a moving-window Mantel analysis was implemented along the HFP gradient. Samples were first ranked in ascending order according to their HFP values. A sliding window with a fixed size of 10 samples was then moved along this gradient with a step size of one sample. Within each window, we calculated the Mantel correlation (*r*) between the environmental distance matrix (spatial and climatic variables and water properties) and the network topological distance matrix (transitivity, average degree, centralization, and assortativity). Second, we assessed the correlations between the Euclidean distance matrices of MMF/PBR and the Bray-Curtis dissimilarity matrices of taxonomic composition and network structure.

### Random forest models

To evaluate the relative importance of biotic and abiotic drivers on both node rewiring scores and network complexity, random forest models were employed using the R rfPermute package^87^. For node-level predictors, the topological baseline (Max_degree) was defined as the maximum degree a specific node achieved across the two regional networks. This metric serves as a quantitative proxy for a taxon’s interaction capacity, reflecting its interaction requirements and fundamental position within the communities. The environmental optima for each ASV (e.g., Opt_salinity and Opt_pH) were mathematically computed as the abundance-weighted mean of the respective environmental variable across all samples where the ASV was present. All random forest models were constructed using 1,000 decision trees (ntree = 1000) with the default mtry parameter to ensure the robust convergence of error estimates. The relative importance of each predictor was quantified by the percentage increase in Mean Squared Error (MSE) upon variable permutation.

To validate whether local topological rewiring can change ecosystem effects, we developed an *in silico* node-centered rewiring simulation framework in Python. The protocol consisted of two phases: graph embedding-based predictive modeling and targeted topological perturbation. First, sample-specific subnetworks were extracted. To translate these discrete topologies into continuous numerical features, each sample subnetwork was embedded into an 18-dimensional latent representation using the Graph2Vec algorithm (Weisfeiler-Lehman iterations = 1; node labels defined as kingdom and phylum). Random forest models were trained on these graph embeddings to predict MMF and potential PBR. Model robustness was rigorously evaluated via 10-times repeated 5-fold cross-validation. During the rewiring simulation, a targeted full-template substitution scheme was applied. At each iteration, a shared focal node was probabilistically selected, weighted by its empirical rewiring score multiplied by the Jaccard mismatch between its current neighborhood and the target template. The existing local edges of this focal node were in silico deleted and seamlessly replaced by the interaction topology observed in the target regional network (strictly restricted to nodes active in the current sample). Crucially, this manipulation altered only the local network structure without modifying node identity or abundance. Following each localized rewiring event, the updated subnetwork was dynamically re-embedded, and the pre-trained random forest models computed the simulated shifts in MMF and PBR. This standardized procedure was executed across representative Tajikistani (T2_1 and T2_4; lower MMF but higher PBR) and Chinese (X9_8 and X9_10; higher MMF but lower PBR) subnetworks, generating 12 independent continuous trajectories (15 rewiring steps each) to map the changes in MMF and PBR.

### Generalized linear modeling

For MMF, we integrated the abundances of functional genes involved in carbon, nitrogen, phosphorus, and sulfur cycles. For PBR, we aggregated the abundances of ARGs, VFs, potential human pathogens, and toxic algae. The raw abundance of each constituent variable was first standardized to a uniform scale ranging from 0 to 1 using the min-max normalization method. The standardized values of the respective variables were then averaged to derive the final MMF and PBR indices for each sample. To quantify the relative influence of environmental, community, and network drivers on MMF and PBR, we constructed two hierarchical GLM frameworks. The first framework (baseline model) incorporated a pool of environmental variables, including spatial and climatic variables, water properties, and HFP, as well as community attributes, including Shannon index and composition (PCoA1 and PCoA2 axes) of both prokaryotes and eukaryotes. The second framework (extended model) built upon this baseline by additionally incorporating network topological variables, including node counts (total, prokaryotic, and eukaryotic), edge numbers (total, intra-kingdom, and cross-kingdom), network complexity, and composition of the top four network modules. Prior to model construction, a collinearity diagnostic was performed independently for Chinese and Tajikistani datasets. Within each specific predictor pool, variables exhibiting strong pairwise Pearson correlations (|*r*| > 0.8) with a variance inflation factor (VIF) > 5 were identified and sequentially removed.

The GLMs were implemented using the h2o R package, employing an Elastic Net regularization approach with the parameters “family = ‘gaussian’, alpha = 0.5, and lambda_search = TRUE”^88^. The elastic net penalty (alpha = 0.5) was strategically selected to balance Lasso (L1) and Ridge (L2) regularizations, which is particularly effective for ecological datasets characterized by high-dimensional predictors and potential multicollinearity. This allowed for robust variable selection while maintaining stability among grouped correlated predictors. The quantitative comparisons of the relative influence of each variable were conducted through standardized model coefficients, and the total contribution of each model was expressed as the percentage of explained variance.

## Notes

### Competing Interest Statement

The authors have declared no competing interest.

### Funding

- National Key R&D Program of China: 2025YFE0103900

